# Compositional and Functional Profiling of the Gut Microbiome in Sarcoidosis

**DOI:** 10.1101/2025.09.23.677246

**Authors:** Jessica M Lee, Yang Chen, Yue Huang, Montserrat Hernandez Martines, Russell Edafetanure-Ibeh, Christen L Vagts, Christian Ascoli, Jun Sun, David L Perkins, Nadera Sweiss, Patricia W Finn

## Abstract

**Background:** Sarcoidosis is a chronic inflammatory disease characterized by systematic granuloma formation, predominantly in the lungs. One leading hypothesis posits that prolonged exposure to microbial antigens may trigger chronic and dysregulated inflammation. Prior studies have examined the lung microbiota of sarcoidosis patients, but the role of the gut microbiota along the gut-immune axis remains largely unexplored. We elucidate the community composition and function of the gut microbiome in sarcoidosis.

**Methods:** Subjects diagnosed with sarcoidosis (n=37) were recruited and matched with healthy control subjects (n=37). Stool samples were collected from sarcoidosis patients for metagenomic sequencing, followed by taxonomic classification and functional annotation as KEGG Orthology (KO) pathways. Differential analysis between sarcoidosis and control cohorts was performed using both standard and compositional approaches.

**Results:** Sarcoidosis microbiomes exhibited similar alpha diversity but significantly different beta diversity from control microbiomes (p<0.05). At the phylum level, the proportion of Actinobacteria and Firmicutes expanded whereas Bacteroides and Verrucomicrobia proportionally shrunk in sarcoidosis gut microbiomes compared to healthy control ones. At the species level, 1,495 species were significantly more abundant and 279 less abundant in sarcoidosis compared to controls (FDR<0.05). Of these, 13 species demonstrated greater intergroup than intragroup variation (effect size>1), including microbes previously detected in the lungs and granulomas of sarcoidosis. Functional profiling revealed 409 KO pathways that were significantly underrepresented in sarcoidosis compared to control (FDR<0.05), of which 79 met an effect size > 1. These include pathways in amino acid and energy metabolism as well as HIF-1 signaling.

**Conclusions:** Our findings highlight the potential role of gut microbiome composition and function in inflammatory processes that are associated with granuloma formation in sarcoidosis and warrant further investigation into gut hyperoxia-driven dysbiosis.

## BACKGROUND

Sarcoidosis is a chronic inflammatory disease characterized by the infiltration of affected organs by immune cell clusters known as granulomas. These granulomas comprise epithelioid cells, mononuclear cells, CD4+ T cells, and some CD8+ T cells that generally form around an offending antigen. They predominantly target the lungs, but they can form in any organ system, including the skin, eyes, intestines, heart, and brain. As a result, sarcoidosis patients often present with non-specific symptoms that overlap with other inflammatory diseases, making it difficult to diagnose and treat^1,2^. Prior studies implicated microbial antigens as a trigger for inflammation driving granuloma formation through activation of Th1 and Th17 cells^3^. However, efforts to identify culpable microbial targets from the the lung microbiome in sarcoidosis were met with inconsistent results^4–6^. Another source of microbial antigens within the body is the gut microbiome, which is known to crosstalk with the lungs through the blood circulatory system, referred to as the gut-lung axis^7^. Moreover, dysbiosis of the gut microbiome has been linked to other chronic pulmonary and inflammatory diseases, though it has yet to be characterized in sarcoidosis^8^. We aimed to address this gap by profiling the heretofore uncharacterized gut microbiome in sarcoidosis patients and highlighting microbial and functional markers of immune pathology along the gut-lung axis.

## METHODS

### Cohort recruitment and sample acquisition

Study approval was obtained through the University of Illinois at Chicago (UIC) IRB Ethics Review Committee, Approval #2016-0063. Subjects with biopsy-proven sarcoidosis, diagnosed in accordance with ATS/ERS/WASOG criteria^9^, were recruited. All subjects were older than 18 years of age and received their sarcoidosis care in the Bernie Mac Sarcoidosis Translational Advanced Research (STAR) Center at UIC. Subjects who had comorbid inflammatory disease that may afflict confounding effects on gut microbiome composition, including malignancy, connective tissue or autoimmune disease, or active infection were excluded. To study the gut microbiome, stool samples were collected between December 2017 and January 2021 and flash frozen at -80⁰C within 2 hours. As a control cohort, healthy gut microbiome data published by Plichta et al. were downloaded from NCBI BioProject and processed using the same bioinformatics pipeline as the sarcoidosis cohort^10^.

Sarcoidosis patient clinical data was also extracted from their medical charts within 30 days of sample collection. Patient severity was defined according to criteria established by our clinical collaborators. Subject demographics included age, sex, self-reported race, body mass index (BMI), and smoking status. Laboratory values included complete blood counts, lymphocyte subsets, serologic markers of inflammation (ESR, CRP), and markers of extrathoracic organ involvement (complete metabolic panels and vitamin D levels). Any patients with chronic sarcoidosis involving neurologic, cardiac, ocular, or advanced pulmonary disease (forced vital capacity (FVC) less than 40% of predicted values) are considered to have severe disease^11^.

### DNA extraction and metagenomic shotgun sequencing

Microbial DNA was extracted from stool using the PowerSoil® DNA Isolation Kit (catalog no. 12888-50) and stored at -80⁰C. DNA concentration was measured using the Tecan SPARK® Multimode Microplate Reader (Tecan, Mannedorf, Switzerland). Purified DNA was fragmented into lengths of 400 bp via sonication, then inserted into libraries using the NEBNext® Ultra™ DNA Library Prep Kit for Illumina kit (catalog no. NEB #E7370L). Library concentration was measured via Qubit fluorometer using the Qubit™ dsDNA High Sensitivity Assay Kit (catalog no. Q32854). Library size and quality was assessed via the Agilent 2100 Bioanalyzer system using the Agilent DNA 7500 Kit (catalog no. 5067-1506). Libraries with sizes ranging between 400 bp and 600 bp and without notable contamination peaks were considered high quality. These high-quality libraries were pooled in 20-sample batches and sequenced on the Illumina MiSeq platform using the Miseq V3 500 kit (catalog no. MS-102-3003), resulting in a sequencing depth of approximately one million reads per sample.

### Taxonomic classification and metabolic pathway annotation

Quality control of the raw demultiplexed reads was performed using *Kneaddata* (v0.10.0) to remove any poor quality or host contaminant reads (https://github.com/biobakery/kneaddata). Briefly, this involved trimming low-quality bases from the 3′ end of reads with *Trimmomatic52* and then discarding trimmed reads <60 nt in length^12^. Host (human) reads were identified and removed by mapping against the human genome (hg19 build) with *Bowtie 2*^13^. Reads passing quality control were then taxonomically classified and quantified as counts using *Kraken2* with *Bracken* as well as *MetaPhlAn* (v4.0.6)^14–16^. Metabolic pathways were also identified based on the previously trimmed reads and mapped to KEGG orthology (KO) terms using *HUMAnN* (v3.7)^17^.

### Exploratory analysis of microbiome diversity

The R packages *Phyloseq*^18^ and *Microbiome*^19^ were used to calculate alpha diversity measures, including observed richness, Shannon index, and Simpson index, as well as beta diversity plots based on principal coordinate analysis (PCoA) using Bray-Curtis dissimilarity. Wilcoxon rank sum tests were performed with FDR correction to assess significant differences between sarcoidosis and control cohorts for each alpha diversity measure. Given the high sparsity (zero inflation) of microbiome count data, a compositional approach using Aitchison distance instead of Bray-Curtis was also performed for the above analyses^20^. Statistical significance of the separation between PCA clusters was assessed using permutational multivariate analysis of variance (PERMANOVA) via the *vegan* package.

Taxonomic and functional profiles of the gut microbiome were generated using *Phyloseq*. For both types of profiling, the respective count data was normalized by total count to obtain relative abundance values. These relative abundances were then aggregated at the phylum level for the taxonomic profiles and to module category and subcategory levels for the functional profiles.

### Differential abundance analysis of taxa and metabolic pathways

Microbial species and metabolic pathways that were differentially abundant between sarcoidosis and control cohorts were identified through a compositional approach using *ALDEx2*^21^ to normalize read counts by central log ratio transformation and model them over a multinomial Dirichlet distribution.

### Enrichment analysis

KO pathways were enriched using the KEGG enrichment tool in the *MicrobiomeProfiler* package^22^. KO terms with FDR-adjusted p < 0.05 were considered significantly enriched.

## RESULTS

### Clinical characterization of sarcoidosis and control cohorts

We initially recruited 50 patients diagnosed with sarcoidosis. These patients were screened for antibiotic treatment and/or ongoing infection, as well as cancer history and chemotherapy, resulting in a final cohort numbering 37. This screened cohort predominantly comprises female and black individuals, both of which are major risk factors of sarcoidosis in the United States^23^, with an age range spanning 28 to 83 years and a mean age of 56 ± 11 years (**Table 1**). Thus, findings derived from this cohort should translate closely to the target patient population for this disease.

**Table 1.**
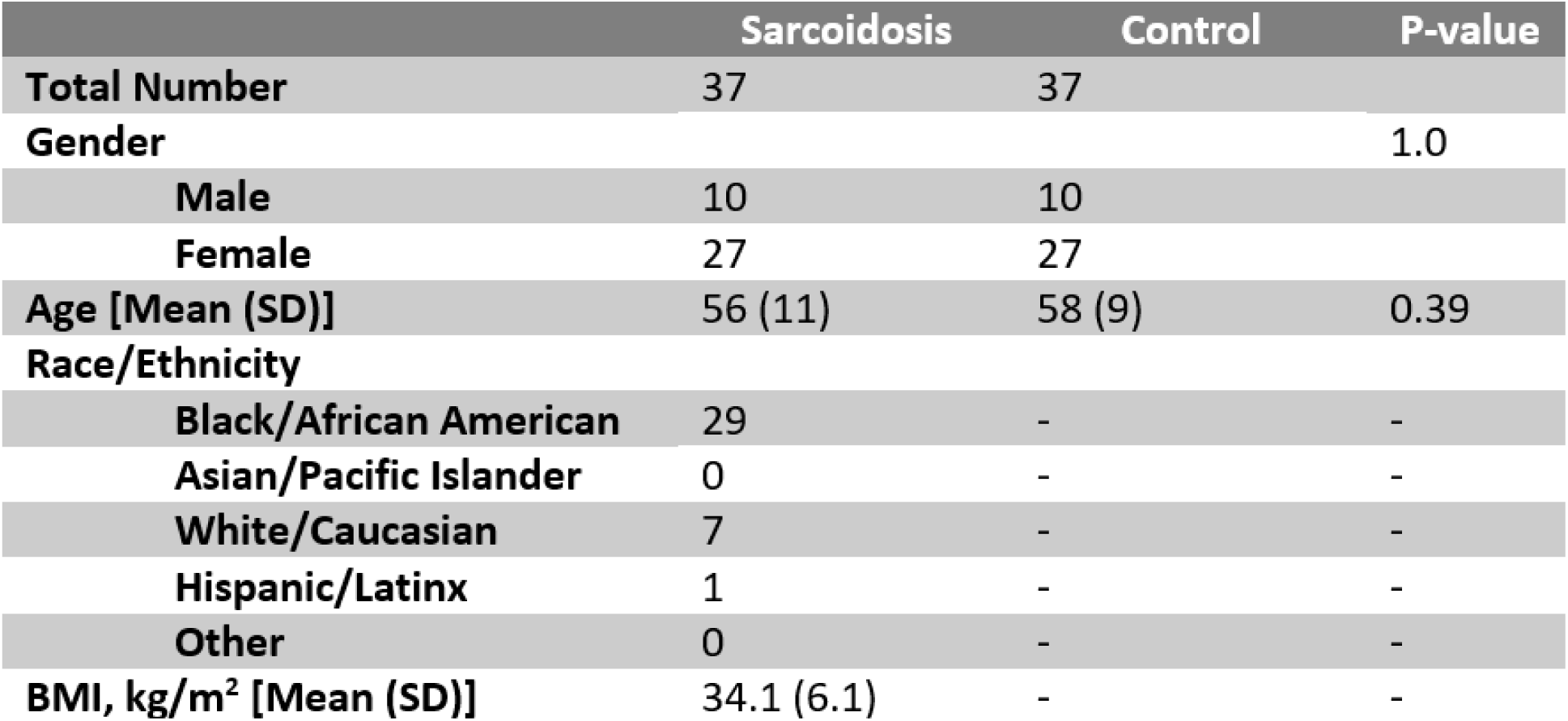
Demographic Data for Sarcoidosis and Control Subjects.

|  | **Sarcoidosis** | **Control** | **P-value** |
| --- | --- | --- | --- |
| **Total Number** | 37 | 37 |  |
| **Gender** |  |  | 1.0 |
| **Male** | 10 | 10 |  |
| **Female** | 27 | 27 |  |
| **Age [Mean (SD)]** | 56 (11) | 58 (9) | 0.39 |
| **Race/Ethnicity** |  |  |  |
| **Black/African American** | 29 | - | - |
| **Asian/Pacific Islander** | 0 | - | - |
| **White/Caucasian** | 7 | - | - |
| **Hispanic/Latinx** | 1 | - | - |
| **Other** | 0 | - | - |
| **BMI, kg/m<sup>2</sup> [Mean (SD)]** | 34.1 (6.1) | - | - |

### Taxonomic profiling of gut microbiome in sarcoidosis

Ever since the advent of metagenomic shotgun sequencing expanded the breadth and depth of microbiome profiling, numerous bioinformatics tools and strategies have been developed to optimize the robustness and completeness of its taxonomic classification^24^. Our chosen bioinformatics pipeline, *Kraken* plus *Bracken*, has been noted to detect more species, including those of low abundance. We began exploring the gut microbiome communities in sarcoidosis and healthy control cohorts through two common measures of community diversity (**Figure 1**). The first, alpha diversity, is a measure of within-sample diversity as defined by richness, the sum total of all species in a community^25^. At the same time, the evenness, or inequality between species abundances, is also important to factor into community diversity, as is done when calculating the Shannon and Simpson indices^26^. Hence, even though the observed richness of the sarcoidosis gut microbiome appears to be significantly (p < 0.05) reduced compared to healthy controls, this reduction was mitigated when accounting for evenness with the Shannon and Simpson indices (**Figure 1A**). Higher richness but comparable evenness of the healthy control gut microbiome relative to sarcoidosis indicates that many of the additional species present may be low in abundance compared to more predominant species, which may remain similarly represented in both cohorts.

**Figure 1.**
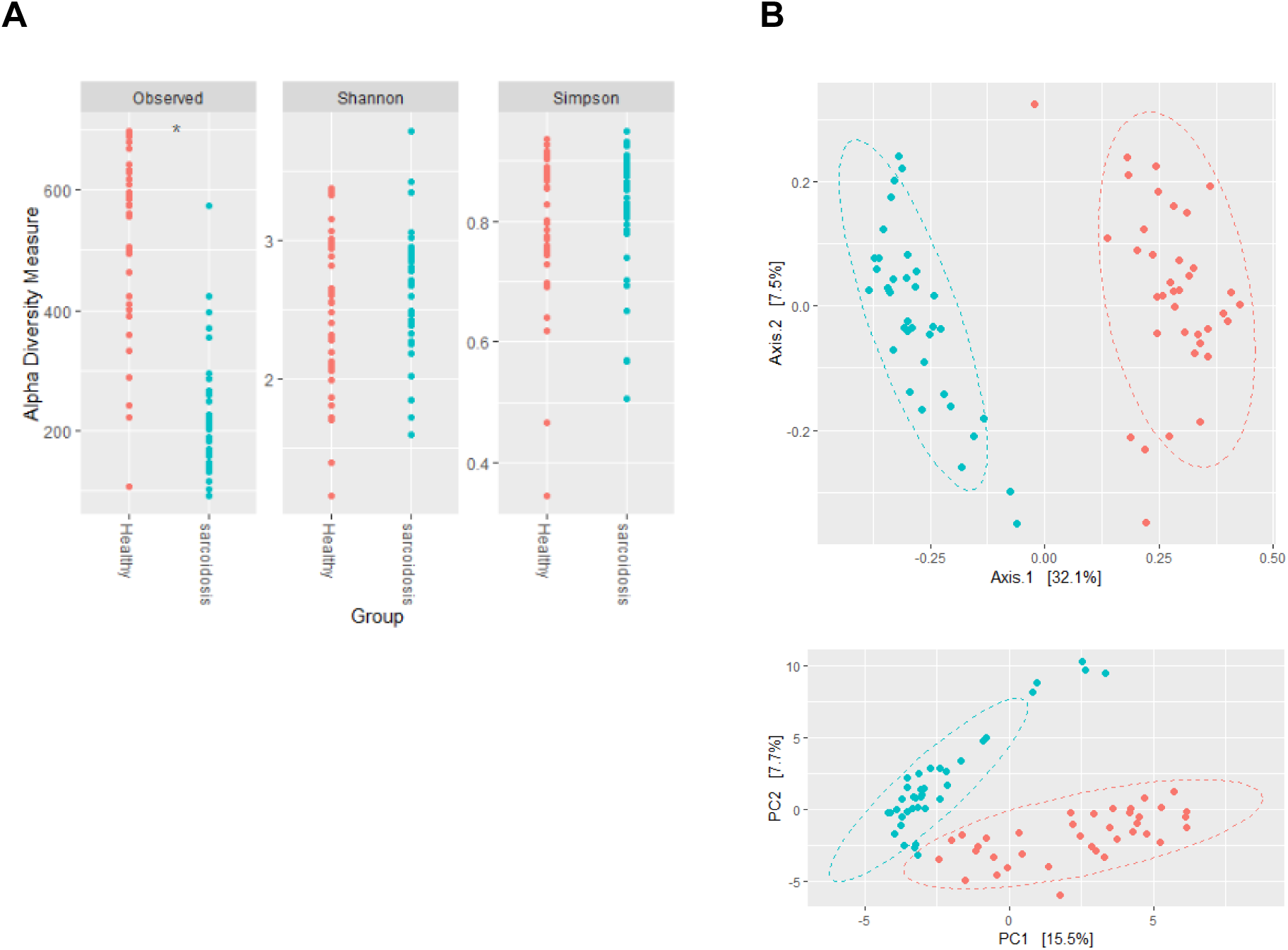
Gut microbiome diversity of healthy control and sarcoidosis cohorts exhibits comparable richness but divergent species membership. (**A**) Alpha diversity measures (observed richness, Shannon index, and Simpson index) are shown for the gut microbiomes in healthy control and sarcoidosis groups based on species count data. (**B**) Principal coordinate analysis (PCoA) plots for beta diversity of the gut microbiomes in healthy control and sarcoidosis groups are shown based on a standard dissimilarity (Bray-Curtis, top panel) or compositional distance (Aitchison, bottom panel) metric. PERMANOVA. *: p≤0.05.

While alpha diversity describes species variability within samples, the other measure of diversity, beta diversity, captures the between-samples species variability^25^. Bray-Curtis dissimilarity is a popular metric based on the number and abundance of shared microbes between communities, but the use of Aitchison distance has gained traction due to its suitability for handling compositional data^20^. Regardless of dissimilarity/distance metric used, we observed a significant (p < 0.001) separation of the sarcoidosis and healthy control clusters, supporting the influence of sarcoidosis on variability in gut microbiome composition between the two groups (**Figure 1B**). Still, while the metric used did not significantly impact the beta diversity verdict, it may be worth noting that the percentage of variation explained by the first two principal components was lower when using Aitchison’s distance. This may in part be due to the sensitivity of Bray-Curtis dissimilarity to shifts in abundance depending on the number of microbes in a community^27^. We likewise performed PCoA on demographic factors such as sex and age but observed no clear clustering based on these factors, indicating that they do not significantly influence microbiome diversity in our cohorts.

To assess how closely the diversity measurements are reflected in the overall microbial composition, we profiled the sarcoidosis and healthy control gut microbiomes at the phylum level (**Figure 2**). In total, 37 unique phyla, including 5,929 species, were detected, with five phyla dominating above the rest: Actinobacteria, Bacteroidetes, Firmicutes, Proteobacteria, and Verrucomicrobia. Of these five, Actinobacteria and Firmicutes comprised higher relative abundances in sarcoidosis compared to in controls. As overabundance of these two phyla had been associated with other inflammatory and autoimmune diseases, this finding may likewise suggest a role for these bacteria in promoting development of sarcoidosis through a pro-inflammatory capacity^8,10^. Moreover, Bacteroidetes and Verrucomicrobia, which comprised lower relative abundances in sarcoidosis, had been linked to anti-inflammatory properties, suggesting microbial involvement in the dysregulation of inflammation that may contribute to sarcoidosis as well^28–30^.

**Figure 2.**
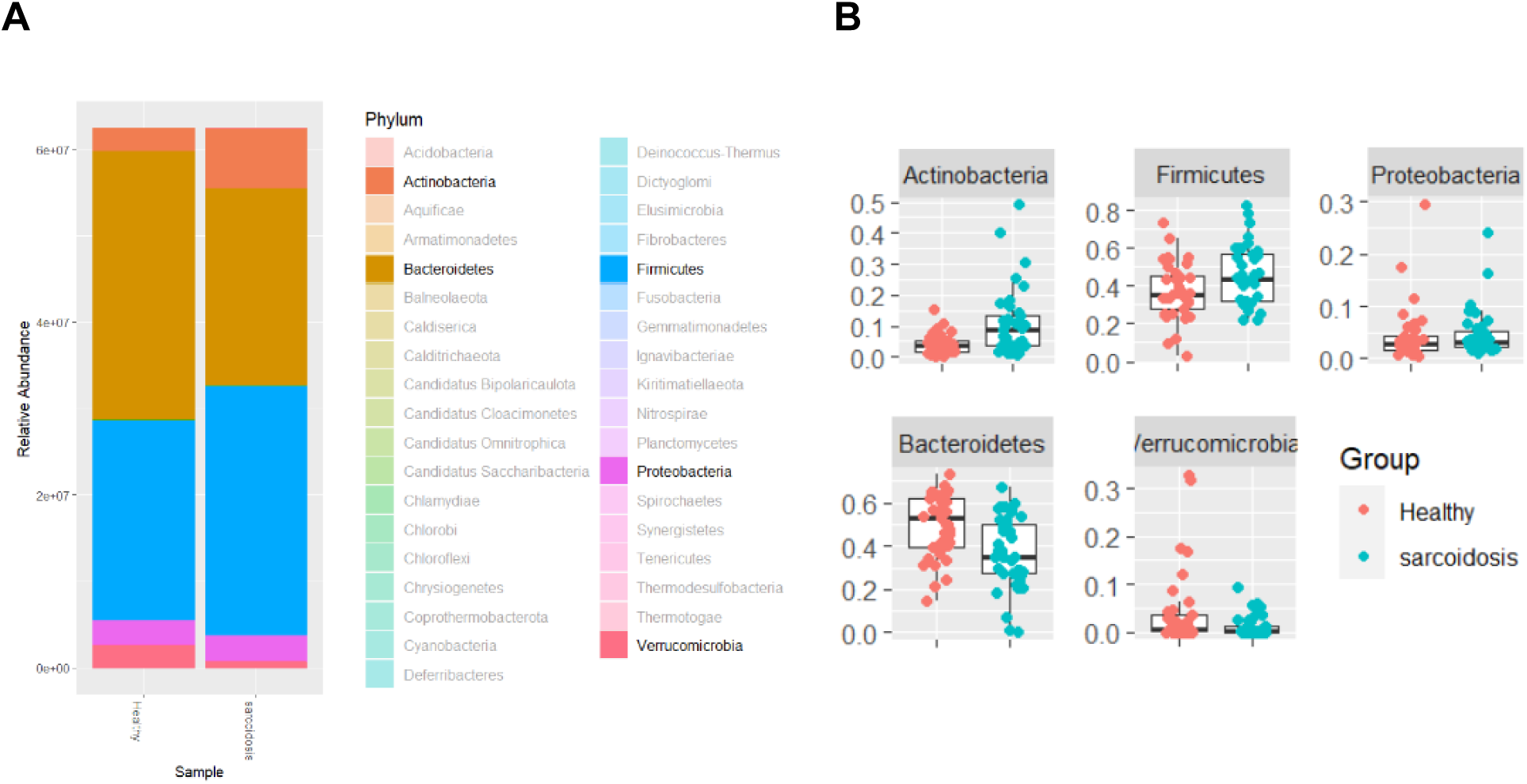
Gut microbiome composition in sarcoidosis patients contains higher proportions of pro-inflammatory phyla and lower proportions of anti-inflammatory phyla compared to healthy controls. (**A**) Gut microbiome composition is represented as stacked bar plots (left) and box-and-whisker plots (right) based on phylum relative abundance. (**B**) The top five most abundant phyla (Actinobacteria, Bacteroidetes, Firmicutes, Proteobacteria, and Verrucomicrobia) are highlighted and individually compared between healthy control and sarcoidosis cohorts, with phyla that are higher in sarcoidosis in the top row and lower in the bottom row.

Having observed differences between sarcoidosis and control gut microbiome profiles at the phylum level, we next investigated which microbial species would be differentially abundant between the two cohorts (**Figure 3**). The standard approach to differential analysis of the microbiome adopted tools from other sequencing modalities, most prominently RNA-seq, which models count data according to a negative binomial distribution. However, the compositional and highly sparse nature of microbiome data has raised concerns over the suitability of these methods, which may exacerbate false positive rates^20,31^. To address these shortcomings, “compositional” approaches have been proposed, such as modeling the count data with a multivariate probability distribution instead^32^. We thus performed differential analysis of the sarcoidosis versus healthy control gut microbiomes using compositional approaches.

**Figure 3.**
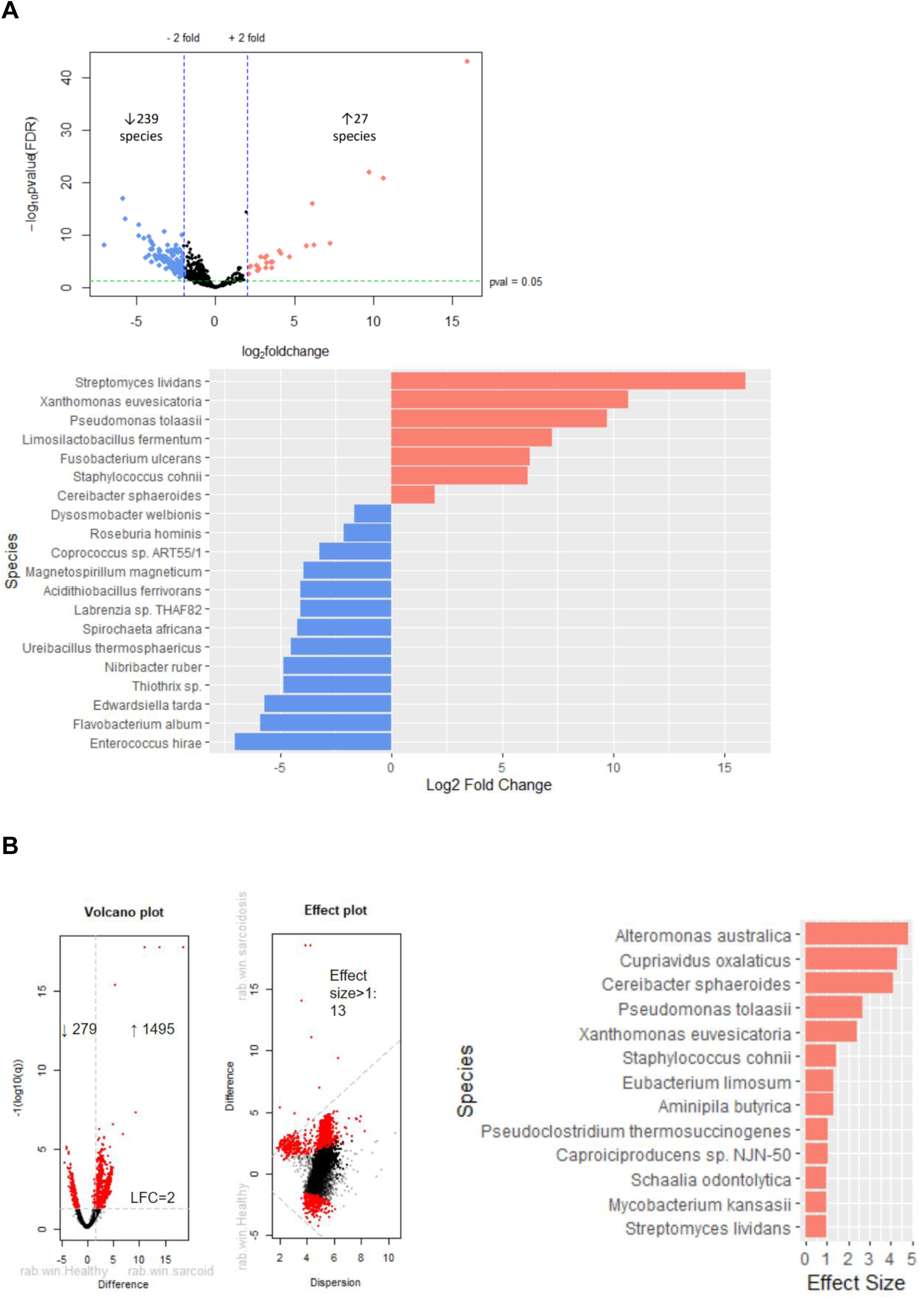
Differentially abundant species in the gut microbiome of sarcoidosis compared to healthy controls present some overlap between standard and compositional approaches. (**A**) Significantly and differentially abundant species in the gut microbiome of sarcoidosis compared to healthy control cohorts were identified using a standard normalization and modeling approach. Volcano plot (top) shows the number of differentially abundant species that are overrepresented (red dots) or underrepresented (blue dots) in the sarcoidosis gut microbiome compared to control based on two criteria: absolute LFC > 2 (black lines) and adjusted p-value (FDR) < 0.05 (green line). The top 20 differentially abundant species in sarcoidosis based on highest absolute LFC and FDR < 0.05 (bottom) are listed in order of greatest increase (red) to greatest decrease (blue). (**B**) Significantly and differentially abundant species in the gut microbiome of sarcoidosis compared to healthy control cohorts were identified using a compositional normalization and modeling approach. Volcano plot (left) shows the number of differentially abundant species that are overrepresented (red dots) or underrepresented (blue dots) in the sarcoidosis gut microbiome compared to control based on three criteria: absolute log2 fold change > 2 (vertical lines) and adjusted p-value (FDR) < 0.05 (horizontal line), as well as absolute effect size >1 (diagonal lines in central effect plot). To the right, the 13 differentially abundant species matching these criteria are listed with the LFC difference within and between the sarcoidosis and healthy control cohorts.

Implementing the standard approach yielded a total of 266 significantly (FDR < 0.05) and differentially (absolute LFC > 2) abundant species, including 27 that increased and 239 that decreased (**Figure 3A**). Of the top 20 most differentially abundant species, *Pseudomonas tolasii* was notable as a microbe previously reported in the blood and lungs to be associated with sarcoidosis^33^. Interestingly, the compositional approach yielded a much greater total of 1,774 species that were significantly and differentially abundant, including 1,495 that increased and 279 that decreased (**Figure 3B**). This likely reflects how standard normalization methods are biased toward finding a small subset of differentially expressed genes^34^. Nonetheless, applying an effect size cutoff of 1 narrowed the number of significant and differentially abundant species to 13. Five of these species (*Cereibacter sphaeroides*, *Pseudomonas tolasii*, *Staphylococcus cohnii*, *Streptomyces lividans*, and *Xanthomonas euvesicatoria*) had also appeared among the top 20 species identified via standard differential analysis, thus strengthening their standing as truly overabundant species in the sarcoidosis gut microbiome.

### Functional profiling of gut microbiome in sarcoidosis

Most microbiome studies to date focus on identifying key microorganisms that could serve as a signature of a disease^35^. However, conflicting findings and the realization that the taxonomic composition of gut microbiomes varied widely across individuals, and even within the same individual at different time points or geographic locations, raises questions as to the scope of their applicability^36–38^. While the exact membership of the gut microbiome may shift, the ecological niches and metabolic functions they fulfill appear to remain constant, only disrupted in cases of dysbiosis^39^. As a result, tracking changes in the metabolic activity of the gut microbiome may serve as a more reliable marker of a disease process.

Through the use of a bioinformatics tool which can predict pathways from metagenomic sequencing data (HUMAnN), we characterized the functional profiles of the gut microbiome in sarcoidosis and healthy control subjects, as illustrated in **Figure 4**. In total, 2,229 metabolic pathways were identified as KEGG orthology (KO) terms belonging to nine module categories and 28 subcategories. Of these modules, amino acid metabolism, carbohydrate metabolism, energy metabolism, metabolism of cofactors and vitamins, and nucleotide metabolism constituted the majority of the microbiome’s metabolic activity (**Figure 4A**). Within these broader categories, more specific subcategories such as ATP synthesis, branched-chain amino acid metabolism, carbon fixation, cofactor and vitamin metabolism, cysteine and methionine metabolism, glycosaminoglycan metabolism, methane metabolism, and purine and pyrimidine metabolism appear to be more active (**Figure 4B**). Most all of these modules share similar relative abundance values between the sarcoidosis and healthy control cohorts, reflecting how, even if the constituent gut microbes differ between the two, the same vital metabolic functions are maintained. At the same time, there were other KO terms not assigned to any module categories or subcategories, leaving open the possibility of differences in metabolic activity between the sarcoidosis and healthy control gut microbiomes in these uncaptured pathways.

**Figure 4.**
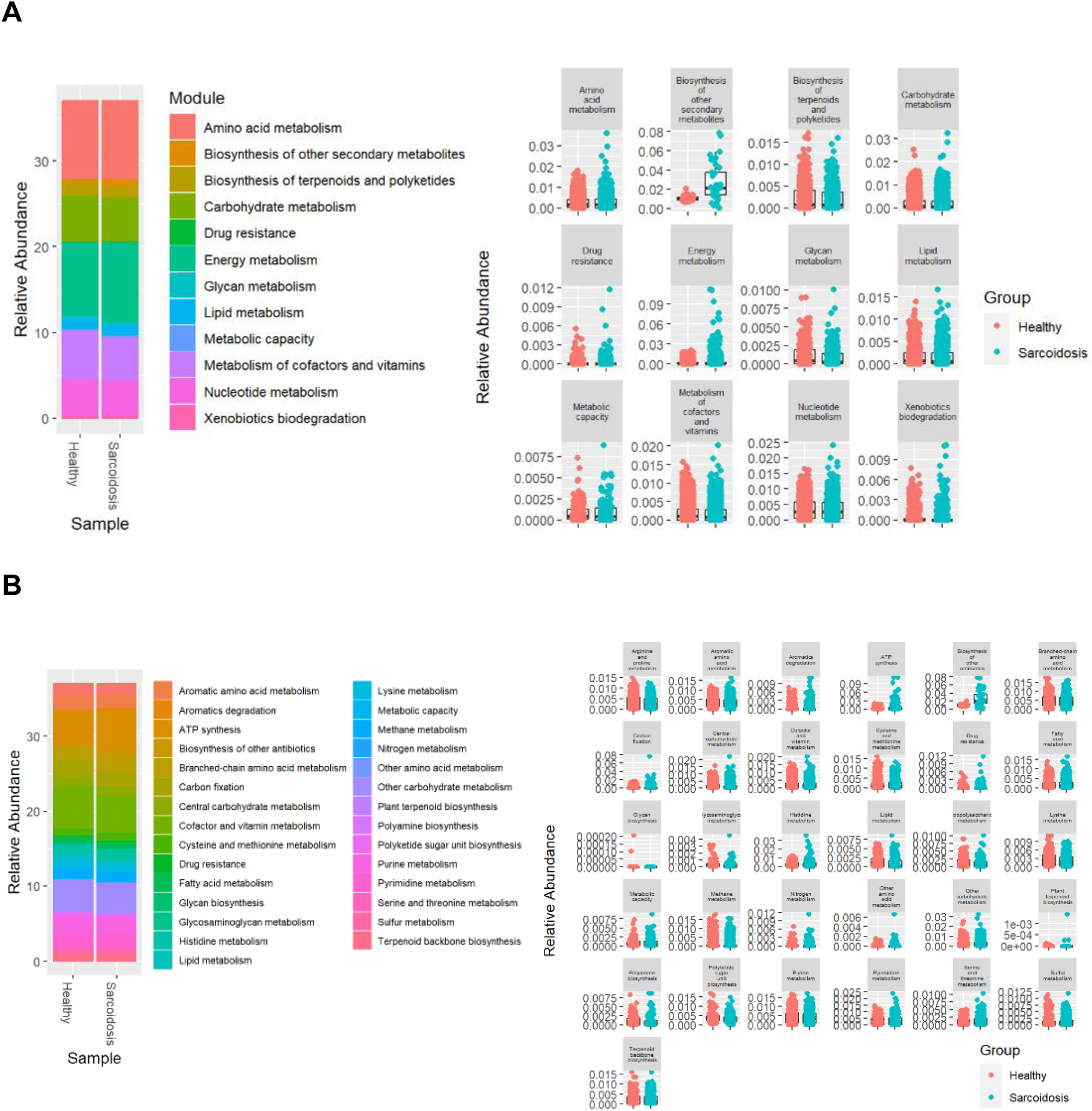
Metabolic pathway modules in sarcoidosis patients compared to in healthy controls. The relative abundance of KEGG orthology (KO) modules in the sarcoidosis and healthy control gut microbiomes are displayed as stacked bar plots (left) and box-and-whisker plots (right). KO modules were identified and compared between healthy control and sarcoidosis cohorts based on (**A**) nine overarching categories and (**B**) 28 subcategories.

While gut microbiome function appeared to be comparable between sarcoidosis and healthy controls at the module level, we investigated if any differences would emerge at the pathway or reaction level. We implemented the same approach for determining differentially abundant species and performed both the standard and compositional versions of differential analysis on KO pathways (**Figure 5**). Standard differential analysis yielded a total of 1,760 pathways, 22 of which were significantly (FDR > 0.05) and differentially (LFC > 2) overrepresented and 23 which were diminished (**Figure 5A**). KEGG enrichment analysis was performed on these select KO terms to reveal significant enrichment (FDR < 0.05) in six metabolic processes, including those related to the ribosome, phosphotransferase system (PTS), carbon metabolism, glyoxylate and dicarboxylate metabolism, starch and sucrose metabolism, and propanoate metabolism. Meanwhile, compositional differential analysis yielded a total of 922 pathways, of which 409 were significantly and differentially underrepresented, and 79 additionally passing the effect size threshold of 1 (**Figure 5B**). KEGG enrichment analysis of these significantly differentially abundant pathways revealed 21 metabolic processes, including those also enriched in the pathways identified by standard differential analysis such as ribosome function and carbon metabolism. Additionally, there was enrichment in processes involved in the biosynthesis of cofactors, glycolysis/citric acid cycle intermediates, and bacterial antibiotics, as well as the HIF-1 signaling pathway, which regulates shifts in energy metabolism strategies in response to changes in oxygen levels^40^. Unlike the results from standard differential analysis, all of the significantly differentially abundant pathways identified by compositional differential analysis were decreased in the sarcoidosis gut microbiome. Upon closer inspection, the KO pathways found to be increased by standard differential analysis were largely related to PTS, which regulates the transport and phosphorylation of carbohydrates and derivatives only in bacteria^41^. This could suggest an enhanced effort by bacterial members of the gut microbiome to acquire more energy sources in response to decreased carbon metabolic activity in the gut environment, but it may also be a consequence of the fewer number of differentially abundant pathways deemed significant by the standard approach, leading to insufficient KO terms for significant enrichment under the terms enriched in the compositional approach. Taking all findings into consideration, the sarcoidosis gut microbiome may fulfill the same metabolic functions as the healthy control gut microbiome but exhibits diminished activity with regards to metabolism or biosynthesis of carbohydrates, amino acids, nucleic acids, cofactors, and bactericidal compounds, which could be related to dysregulated energy metabolism responses to stress conditions such as nutritional scarcity and hyperoxia.

**Figure 5.**
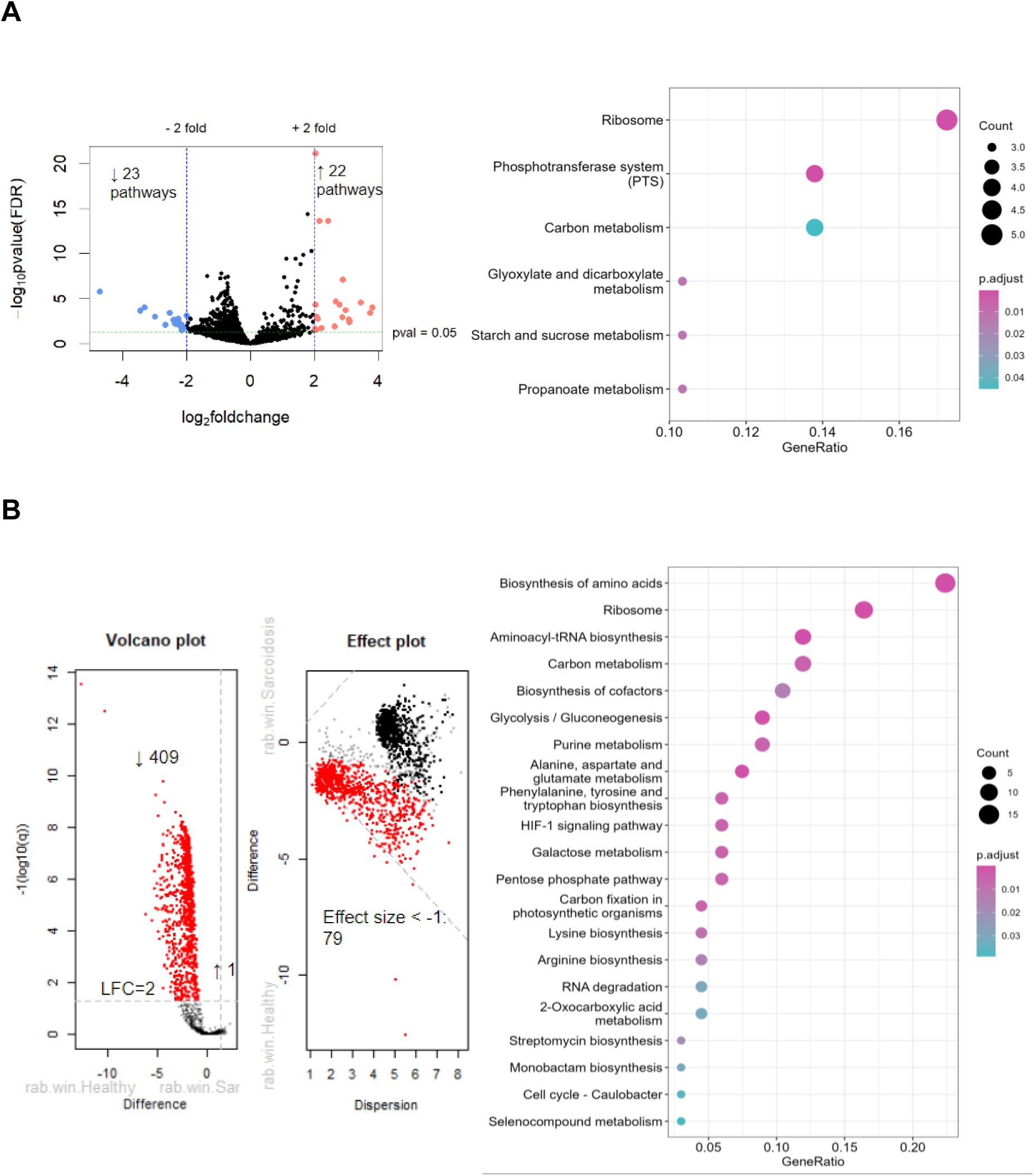
Differentially abundant metabolic pathways in the gut microbiome of sarcoidosis and controls. (**A**) Significantly and differentially abundant metabolic pathways representative of gut microbiome functions in sarcoidosis compared to healthy control cohorts were identified using a standard normalization and modeling approach. Volcano plot (left) shows the number of differentially abundant pathways that are overrepresented (red dots) or underrepresented (blue dots) in the sarcoidosis gut microbiome compared to control based on two criteria: absolute LFC > 2 (black lines) and adjusted p-value (FDR) < 0.05 (green line). Enrichment analysis of these differentially abundant pathways is displayed to the right. (**B**) Significantly and differentially abundant pathways in the gut microbiome of sarcoidosis compared to healthy control cohorts were identified using a compositional normalization and modeling method. Volcano plot (left) shows the number of differentially abundant pathways that are overrepresented (red dots) or underrepresented (blue dots) in the sarcoidosis gut microbiome compared to control based on three criteria: absolute LFC > 2 (vertical lines) and adjusted p-value (FDR) < 0.05 (horizontal line), as well as absolute effect size >1 (diagonal lines in central effect plot). Enrichment analysis of these differentially abundant pathways is displayed to the right.

## DISCUSSION

We present the taxonomic and functional profiles of the gut microbiome in sarcoidosis as compared to healthy control subjects. By comparing the diversity and differentially abundant species and pathways identified by standard versus compositional approaches, we can evaluate our results with a more critical lens to make informed conclusions about microbiome communities and functions. In terms of alpha diversity, the sarcoidosis gut microbiome exhibited lower observed richness of species compared to the healthy control gut microbiome, but this difference became less significant when accounting for the evenness of species distribution, suggesting the discrepancy in richness is attributed to presence of low-abundance species in the healthy control gut microbiome (**Figure 1A**). Moreover, PCoA of the member species to determine beta diversity revealed significant separation between the sarcoidosis and healthy control clusters, indicating that the gut microbiomes of sarcoidosis patients are compositionally distinct from those of healthy control subjects (**Figure 1B**). We also considered the possibility that this variability could be driven by differences in the site of sample collection and processing for our sarcoidosis versus control cohorts, and PCoA was performed on our cohorts with gut microbiome datasets of other cohorts from the same and different geographic regions. While the cohorts did not show clear clustering according to geographic region, we found that their clustering more closely aligned according to the bioinformatics tool that was used for taxonomic classification. Since our control cohort was analyzed using the same bioinformatics pipelines as our sarcoidosis cohort, this confirmed that the observed variability between sarcoidosis and control gut microbiomes should be driven by intrinsic differences of the two cohorts.

In accordance with these diversity observations, we noted that both sarcoidosis and control gut microbiomes were predominantly composed of the same five phyla (Actinobacteria, Bacteroidetes, Firmicutes, Proteobacteria, and Verrucomicrobia) but with different proportions relative to the rest of the microbiome (**Figure 2**). Actinobacteria and Firmicutes were demonstrably increased while Bacteroidetes and Verrucomicrobia were decreased in the sarcoidosis compared to control gut microbiome. Proportions of Proteobacteria appeared comparable between sarcoidosis and controls, but they are known to be pro-inflammatory due to the lipopolysaccharides (LPS) on their cell surface which trigger inflammation through activating macrophages^42^ and have been found to be enriched in the gut microbiome for other inflammatory and autoimmune diseases^8,43^. Interestingly, Actinobacteria and Proteobacteria were also found to be significantly more abundant in the lung microbiome of sarcoidosis patients, suggesting potential interactions via the gut-lung axis^44^.

To better evaluate the differentially abundant species and pathways for significance, we compared the results of the standard versus compositional approaches. While the compositional approach identified 1,774 species as significantly (p < 0.05) and differentially (LFC > 2) abundant, we additionally implemented an effect size cut-off of 1, which filters out species with greater variation within each cohort than between cohorts, resulting in a highly confident set of 13 differentially abundant species (**Figure 3**). From these 13, five were also identified by the standard approach: *Cereibacter sphaeroides, Pseudomonas tolasii, Staphylococcus cohnii, Streptomyces lividans,* and *Xanthomonas euvesicatoria*.

While these species are differentially abundant in our sarcoidosis cohort, they may not necessarily apply broadly to all patients with sarcoidosis due to the wide variability in gut microbiome composition among individuals living in different geographic regions and environments^45^. In contrast, the functional profile of the human gut microbiome would remain more consistent across individuals, even if such functions are carried out by different microbes^39^. Therefore, we also sought to identify which metabolic functions may be disrupted in the sarcoidosis gut microbiome. Overall, the functions, represented as KEGG module categories and subcategories, and metabolic activity, measured as relative abundance of each KEGG module, were comparable between sarcoidosis and healthy control subjects (**Figure 4**). Nonetheless, within each module, specific metabolic pathways and reactions may still vary between the sarcoidosis and control gut microbiomes at the KO pathway level. Differential analysis of the KO pathways revealed twice as many differentially abundant pathways identified by the compositional as by the standard approach, and KEGG enrichment of these pathways revealed an even larger discrepancy, with “ribosome” and “carbon metabolism” as the only concordant processes between the two approaches (**Figure 5**). The enrichment terms for the standard approach may be overly broad due to too few KO pathways deemed significant enough to enrich for more specific KEGG terms as they did for the compositional approach. The superior performance of the compositional approach over the standard approach in identifying both differentially abundant species and pathways thus corroborates its suitability for analyzing microbiome data. Among the KEGG terms enriched for this approach, many are related to energy metabolism, from mediators of the glycolysis and the citric acid cycle to synthesis of their intermediates. The reduced activity of these functions and the HIF-1 signaling pathway, coupled with the enhanced activity of PTS as indicated by standard differential analysis, may be contextualized by the synergistic relationship between commensal microbes and intestinal epithelial cells (IECs), in which IECs take up microbial metabolites produced via fermentation to generate energy through consuming oxygen via mitochondrial oxidative phosphorylation, thus maintaining a hypoxic environment for continued fermentation by these microbes^46–47^. Disruption of this process, whether by insufficient substrates for fermentation or elevation in oxygen levels, would lead to further exacerbating nutrient starvation and intestinal hyperoxia. Hyperoxia in the lung and gut microbiomes has been associated with increased abundance of the aerobic phyla Proteobacteria and Actinobacteria and decreased abundance of anaerobes like Firmicutes and Bacteroidetes^48–49^. While the taxonomic profile we characterized for the sarcoidosis gut microbiome featured an increase instead of decrease in Firmicutes, these may comprise facultative anaerobes like the differentially abundant species *Staphylococcus cohnii* which expanded in place of their obligatory anaerobic counterparts. As HIF-1 also regulates IEC barrier function, its downregulation may compromise the integrity of the epithelial barrier, potentially leaking LPS and other inflammatory microbial components into the bloodstream^46^. Notably, HIF-1 is also highly expressed by sarcoid macrophages and granulomas and implicated in sarcoidosis pathogenesis through regulating Th17 differentiation as well as the macrophage proinflammatory response during hypoxia^50–51^. In addition to hypoxia, HIF-1 is activated by iron deprivation, as iron is required for its degradation by prolyl hydroxylases during normoxic conditions^52^. To ensure their own nutritional needs, gut microbes are known to produce metabolites that suppress HIF-2α regulation of host iron transport, and thus may contribute to overexpression of HIF-1 in sarcoidosis by driving a state of hypoxia and/or iron deprivation^53^. The question of which means may trigger HIF-1 overexpression in sarcoidosis can be addressed by referring to any iron tests and blood gas tests performed on our sarcoidosis subjects.

A limitation to our functional analysis is the incompleteness of our KEGG module mapping and enrichment analysis methods, as the subset of KO pathways that failed to be mapped/enriched could pertain to valuable functions that would not have been captured in our functional profiling. One method to address this shortcoming is manually curating module assignments for all KO pathways and then performing gene set enrichment analysis to incorporate information on their significance and differential abundance into determining the most significantly altered modules^54^. Moreover, though our study is sufficiently powered according to power analysis calculations, increasing our sample size could be beneficial toward studying subgroups within the sarcoidosis cohort to assess the impact of certain clinical features, such as sarcoidosis severity, medication types, and comorbidities, on variation between gut microbiome profiles within the larger cohort.

Our study characterized the gut microbiome community and function in sarcoidosis, revealing insights that can guide further understanding of sarcoidosis etiology and expand diagnostic and therapeutic options with targeting the gut microbiome.

